# End-to-end simulation of nanopore sequencing signals with feed-forward transformers

**DOI:** 10.1101/2024.08.12.607296

**Authors:** Denis Beslic, Martin Kucklick, Susanne Engelmann, Stephan Fuchs, Bernhard Y. Renard, Nils Körber

## Abstract

**Motivation:** Nanopore sequencing represents a significant advancement in genomics, enabling direct long-read DNA sequencing at the single-molecule level. Accurate simulation of nanopore sequencing signals from nucleotide sequences is crucial for method development and for complementing experimental data. Most existing approaches rely on predefined statistical models, which may not adequately capture the properties of experimental signal data. Furthermore, these simulators were developed for earlier versions of nanopore chemistry, which limits their applicability and adaptability to the latest flow cell data.

**Results:** To enhance the quality of artificial signals, we introduce *seq2squiggle*, a novel transformer-based, non-autoregressive model designed to generate nanopore sequencing signals from nucleotide sequences. Unlike existing simulators that rely on static k-mer models, our approach learns sequential contextual information from the signal data. We benchmark *seq2squiggle* against state-of-the-art simulators on real experimental R10.4.1 data, evaluating signal similarity, basecalling accuracy, and variant detection rates. *Seq2squiggle* consistently outperforms existing tools across multiple datasets, demonstrating superior similarity to real data and offering a robust solution for simulating nanopore sequencing signals with the latest flow cell generation.

**Availability and Implementation:** *seq2squiggle* is freely available on GitHub at: github.com/ZKI-PH-ImageAnalysis/seq2squiggle

## 1 Introduction

Long-read nanopore sequencing has emerged as a transformative technology in the field of genomics, offering rapid and cost-effective DNA sequencing capabilities with applications ranging from fundamental research to clinical diagnostics (Giesselmann, 2021). The translocation of analyte molecules through a nanometer-sized pore generates a distinct current signal that represents the physical properties of the molecule inside the pore. These signals are then translated into corresponding DNA sequences (Delahaye and Nicolas, 2021). Nanopore sequencing has significantly advanced our ability to detect single nucleotide variations (SNVs) and insertions/deletions (indels), which are crucial for understanding genetic diversity and susceptibility to diseases. SNVs, or single nucleotide polymorphisms, play a pivotal role in genomic diversity and disease susceptibility, while indels involve mutations where nucleotides are either inserted or deleted from the DNA sequence. Accurate detection of these variations is particularly challenging with nanopore sequencing due to potential errors introduced by homopolymer regions, which can lead to false-positive calls (Delahaye and Nicolas, 2021; Wang *et al*., 2021). The potential of long-read sequencing has sparked the development of a large variety of methods for basecalling, modification detection, error correction, genome assembly, and detection of structural variations (Amarasinghe *et al*., 2020).

Simulation of sequencing signals from nucleotide data is crucial to complement experimental nanopore data and to benchmark recently developed methods. It allows researchers to refine experimental protocols, evaluate sequencing performance, and deepen the understanding of the interactions between DNA molecules and nanopores (Li *et al*., 2018). Sequencing simulators such as DeepSimulator (Li *et al*., 2018), NanosigSim (Chen *et al*., 2020), and squigulator (Gamaarachchi *et al*., 2023) generate nanopore sequencing signals using input nucleotide sequences and pre-existing k-mer models. These simulators first calculate the event level of each k-mer based on pore models provided by Oxford Nanopore Technologies (ONT), then sample the duration of each event from a random distribution (e.g., gamma distribution), and finally add Gaussian noise to the signal (Figure 1A). Although DeepSimulator and NanosigSim incorporate deep learning techniques in their methods, these are limited to specific modules for improvements in pore model accuracy or noise generation. Moreover, these simulators were developed and optimized using the earlier R9.4 chemistry and have not been evaluated with the most recent R10.4.1 chemistry, which exhibits a distinct signal profile due to its modified protein pore and two measurement points (Ahsan *et al*., 2024). Of the mentioned tools only squigulator is capable of generating data for the latest pore chemistry.

**Figure 1.**
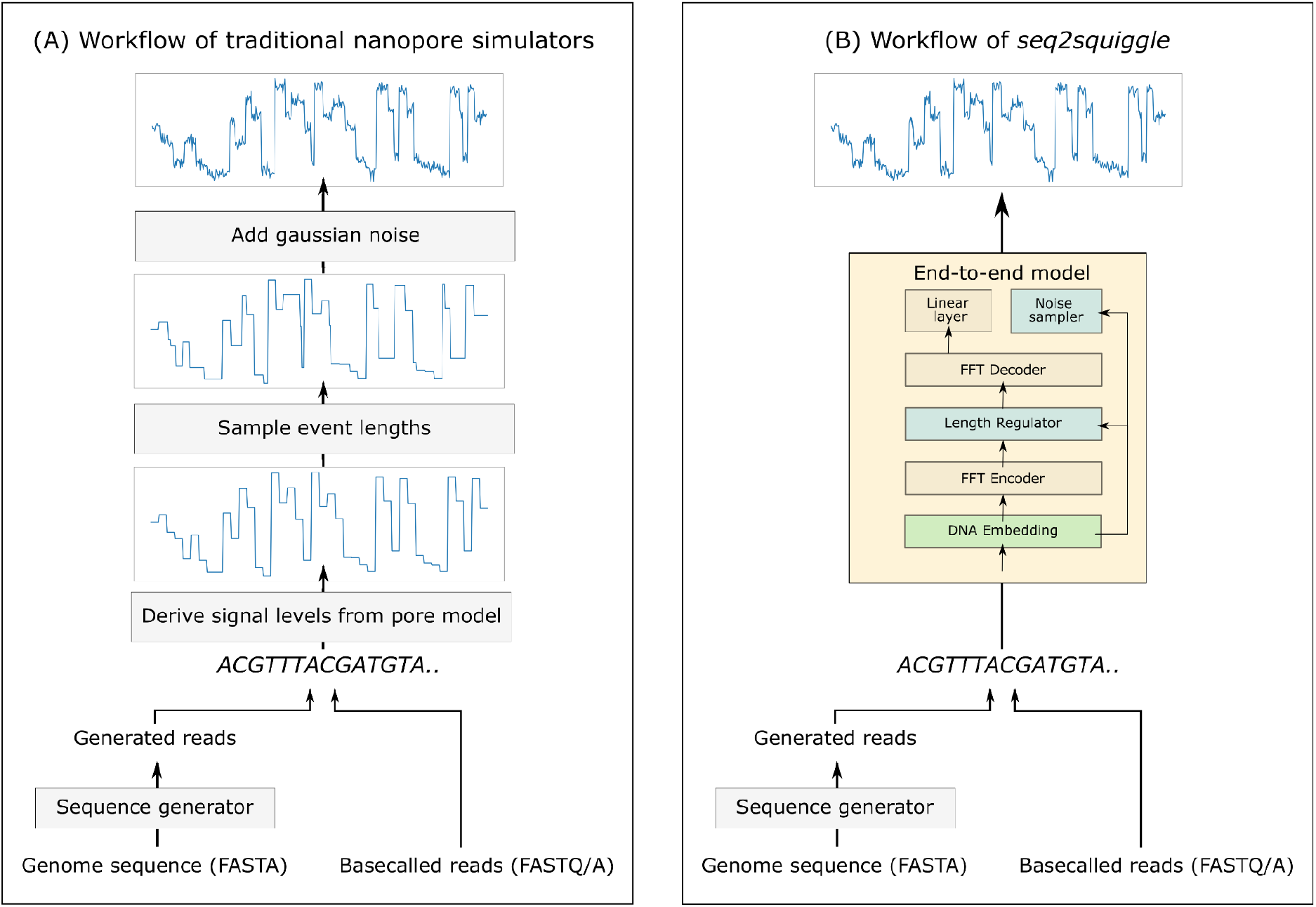
Comparison of nanopore signal simulators with *seq2squiggle*. (A) Traditional nanopore simulators process either read-input (FASTQ/A) or genome-input (FASTA). For genome-input, these simulators first use a sequence generator to produce reads. They then calculate event levels using pre-defined pore models, sample event durations from random distributions, and add Gaussian noise with fixed parameters across all input sequences. While some tools incorporate deep learning for specific sub-modules (e.g., pore models or low-pass filter), these methods are limited to enhancing certain components. (B) In contrast, *seq2squiggle* employs an end-to-end deep learning approach to predict signals directly from nucleotide input, allowing the model to learn signal levels, event durations, and noise distributions from the training data itself.

The reliance on pre-defined pore models and constant statistical assumptions across all sequences poses a risk of inaccuracies in generating in-silico signal data. Teng and coworkers highlighted that the segmentation of raw signals is a critical source of errors in basecalling algorithms (Teng *et al*., 2018), which has led to a paradigm shift in basecalling methodologies towards ‘end-to-end’ deep learning architectures. Applying a similar end-to-end approach to signal simulation offers the potential to enhance the quality of simulated data. These frameworks bypass the segmentation step and directly infer nucleotide sequences from raw signals, improving accuracy and robustness. However, simulating signals from DNA sequences presents a one-to-many mapping challenge, where each DNA sequence corresponds to a variety of possible target signals of alternative event lengths and noise levels. This inherent complexity introduces unique hurdles in defining appropriate loss functions, designing model architectures capable of capturing intricate relationships between sequences and signals, and identifying suitable evaluation metrics. Overcoming these challenges with deep learning frameworks is crucial for improving the accuracy of artificial nanopore signals and supporting the development and benchmarking of new analytical methods.

In this study, we introduce *seq2squiggle*, an innovative transformer-based simulator designed for nanopore sequencing data. Similar to the trend in basecallers, our objective is to predict raw signals from sequence data without directly relying on pore models. By leveraging feed-forward transformer blocks, our model effectively captures broader sequential contexts, enabling the generation of artificial signals that closely resemble experimental observations. Furthermore, *seq2squiggle* includes modules capable of learning event length and amplitude noise distributions from training data (Figure 1B). Evaluation against multiple experimental R10.4.1 datasets reveals that our simulated datasets exhibit a closer resemblance to real-world observations compared to existing tools.

## 2 Methods

### 2.1 General workflow overview

In this section, we present a general overview of our proposed simulator *seq2squiggle*, a novel tool designed for simulating nanopore sequencing signals using feed-forward transformer blocks. Similar to previous simulators, the initial module of our tool is the sequence generator. Given a user-specified reference genome, as well as the desired coverage or number of reads, this module randomly selects starting positions on the genome or contigs to generate sequences that meet the coverage requirements and mimic the length distribution of experimental nanopore reads. *seq2squiggle* can generate reads from a genome to simulate signals or use experimental reads via the --read-input command. The read length distribution can be influenced by various factors, such as the sample species and experimental setup. (Baker *et al*., 2016; Li *et al*., 2018). Given that realistic read-length simulation is complex and not the primary focus of our research, we chose to adopt the read-length distributions previously defined and used in DeepSimulator (Li *et al*., 2018). These include an exponential distribution, a beta distribution, and a mixed gamma distribution, which provide a practical and established framework for our simulator.

To enhance the quality of the simulated signals, *seq2squiggle* avoids the direct use of pore models and instead learns the signal characteristics from data. The details of the signal generation using feed-forward transformers are described in the next chapter.

The simulated reads are either exported to the community-driven SLOW5 format (Gamaarachchi *et al*., 2022) or the new POD5 format by ONT, enabling seamless integration for basecalling and subsequent analysis. This ensures compatibility with existing nanopore sequencing analysis pipelines and supports comprehensive downstream evaluations.

### 2.2 Model architecture

Our proposed model *seq2squiggle* predicts nanopore sequencing signals using a feed-forward transformer (Figure 2A), a deep learning architecture developed and successfully applied for speech synthesis (Ren *et al*., 2019, 2022). Similar to text-to-speech approaches, we map the shorter source sequence (DNA) to the larger target sequence (nanopore signal) for which we employed a feed-forward architecture instead of an autoregressive network.

**Figure 2.**
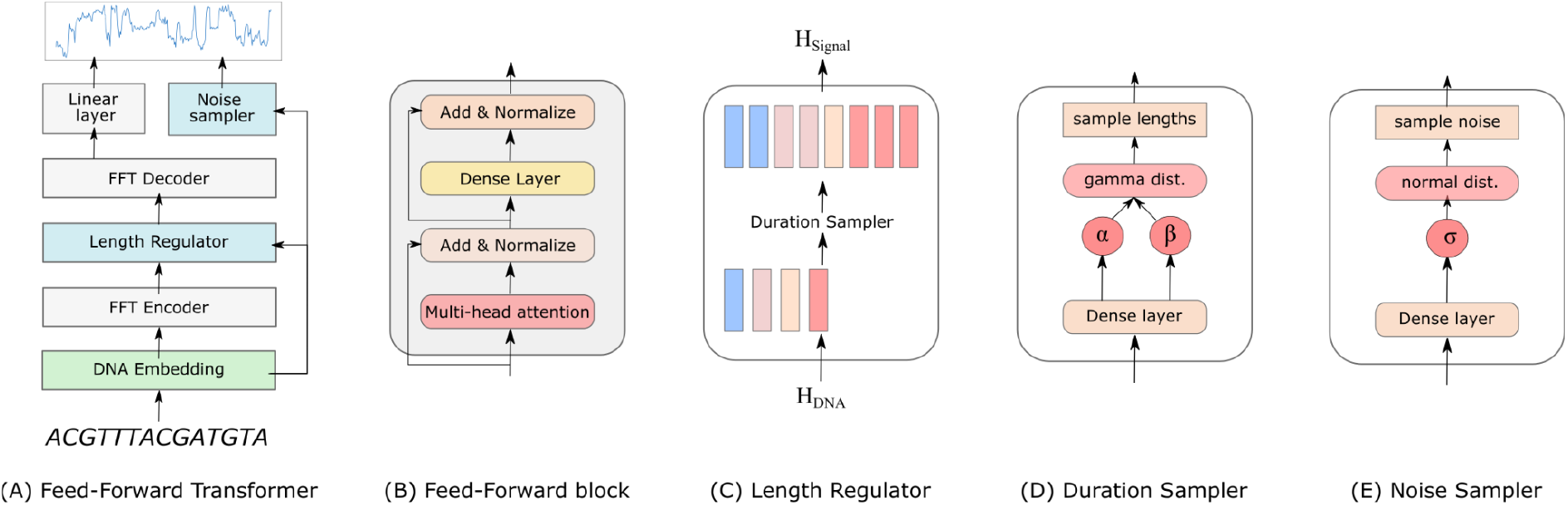
The architecture of *seq2squiggle*. (A) The Feed-Forward Transformer is the core model architecture, using feed-forward networks to map DNA sequences to nanopore signals. (B) The Feed-Foward block consists of multi-head attention and dense layers for cross-position information extraction. (C) The length regulator adjusts the hidden states of the DNA sequences to match the predicted event duration of the nanopore signal. (D) The duration sampler uses a gamma distribution to sample the event lengths based on the DNA embeddings. (E) The noise sampler predicts the standard deviation of gaussian noise based on the DNA embeddings and adds it to the signal to create realistic noise patterns.

The Feed-Forward Transformer (FFT) includes multiple FFT blocks for the DNA to signal transformation, with two blocks on the nucleotide sequence side and two blocks on the nanopore signal side. A FFT block consists of a multi-head attention network and dense layers to extract cross-position information (Figure 2B).

To address the alignment issue between input and target sequences, a length regulator is implemented. This component upscales the hidden states of the DNA sequence according to the event duration of the corresponding nanopore signal (Figure 2C).

To infer the duration lengths of each signal, we employ a duration sampler consisting of two dense layers with ReLU activation resulting in two scalars α and β (Figure 2D). These two scalars are used for parameterizing a gamma distribution from which we sample the duration lengths. Our proposed duration sampler takes the hidden states of the nucleotide sequence into consideration for generating event lengths based on a random distribution. The event lengths from the duration sampler are only used in the inference phase since we can use the reference event duration during training.

Additionally, we incorporate a noise module consisting of two dense layers with ReLU activation, which predicts the standard deviation σ of the signal. Similar to the duration sampler, the prediction is based on the hidden states of the nucleotide sequence, serving as input for the noise module. The predicted standard deviation is then used to sample Gaussian noise distribution, which is subsequently added to the signal after the decoder layer, contributing to the generation of realistic noise patterns (Figure 2E).

Hence, *seq2squiggle* incorporates three loss functions:

1. Signal loss, measured by the mean squared error (MSE), compares the ground truth signal *s*_*ref*_ with the predicted signal *s*_*pred*_.

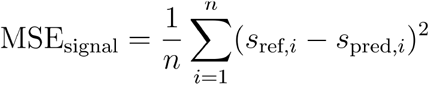
2. Duration loss utilizes the negative log-likelihood (NLL) to observe the reference event lengths *l*_*data*_ given the parameterized gamma distribution with predicted shape θ and scale μ. To ensure that the scale of this loss is comparable to the other loss functions, it is adjusted by the factor τ = 0.0005.

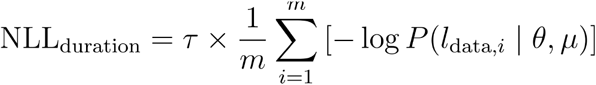
3. Noise loss, also measured by the MSE, compares the standard deviation of the real data *σ*_*ref*_ with the predicted standard deviation *σ*_*pred*_ from the noise module, thereby capturing the variability in the real data.

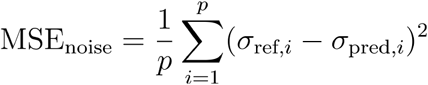

To optimize the model, the three loss functions are combined into a total loss via weighted summation. Summing multiple loss functions, a common approach in multi-task learning (Liang and Zhang, 2020), ensures a balanced optimization process by integrating different aspects of model performance, thereby allowing all objectives to be addressed simultaneously during training:

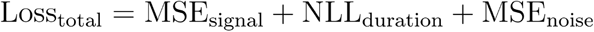

### 2.3 Experimental setup

#### 2.3.1 Datasets for training and evaluation

For our experiments, we used two publicly available R10.4.1 DNA datasets: the Human HCT116 and *Drosophila melanogaster* datasets. Both were sequenced using Kit14 chemistry with a 4kHz sampling rate (Table 1). The raw sequencing files were converted to the SLOW5 format and basecalled using the dna_r10.4.1_e8.2_400bps_sup.cfg model with buttery-eel (v0.4.2) (Samarakoon *et al*., 2023) and the guppy basecall server (v7.1.1). Following basecalling, the reads were aligned with minimap2 (v2.26). The Human HCT116 data was aligned to the hg38 reference genome without alternate contigs, while the *D. melanogaster* data was aligned to Genome Assembly Release 6 plus ISO1 MT, as annotated by the FlyBase Consortium (Jenkins *et al*., 2022). To segment the reads and establish a mapping between the DNA sequence and the nanopore signal, we used the *eventalign* option from uncalled4 (v4.1), a toolkit for nanopore signal alignment developed by Kovaka and coworkers (Kovaka *et al*., 2024).

**Table 1.** Overview of datasets for training and evaluation. Displayed are the name of the dataset, the number of reads, flow cell chemistry, the Accession ID and the corresponding website of each dataset.

| Dataset name | # of total reads | Chemistry | Accession ID | Website |
| --- | --- | --- | --- | --- |
| Drosophila melanogaster | 229,767 | R10.4.1 Kit 14 (4kHz) | PRJNA914057 | https://labs.epi2me.io/open-data-drosophila/ |
| Non-methylated HCT116 | 5,149,102 | R10.4.1 Kit 14 (4kHz) | PRJEB64592 | https://hasindu2008.github.io/f5c/docs/r10train |

For training *seq2squiggle*, we used 160,936,708 chunks derived from 1,417,538 reads of chromosome 2, 3, and 4 (chr2-4) of the Human HCT116 dataset. For validation, we used 100,000 chunks from 56 reads of chromosome 22 (chr22) from the same dataset. For evaluation, we used 100,000 reads from chromosome 1 (chr1) of the Human HCT116 dataset and 100,000 reads from the *D*.*melanogaster* dataset. This approach allowed us to comprehensively assess the performance of *seq2squiggle* across different genomic contexts.

#### 2.3.2 Model configuration and training procedure

*Seq2squiggle* was trained using chunks from the uncalled4 *eventalign* table, divided into segments of 16 overlapping 9-mers and 250 corresponding signal points. Shorter signals were zero-padded, while chunk pairs with signals over 250 signal points were filtered out. The model vocabulary included 5 symbols: the four DNA bases (‘A’, ‘C’, ‘G’, ‘T’) and an empty symbol (‘_’).

Our model architecture features two FFT blocks on both the DNA-encoder and signal-decoder sides, with hyperparameters set as follows: DNA embeddings, self-attention hidden size, and other hidden dimensions all set to 64; feed-forward projection size set to 256; and 8 attention heads (Supplementary Table 1).

The model was trained on 8 NVIDIA A100 GPUs with a batch size of 512 DNA-signal chunks. We utilized the Adam optimizer with parameters β1 = 0.9, β2 = 0.98, and ε = 10e−8. A dropout rate of 0.1 and gradient clipping-by-norm set to 1.0 were applied. The learning rate schedule included a linear warm-up (1% ratio) followed by cosine decay, with a maximum learning rate of 0.00025. The model was trained for 20 epochs.

#### 2.3.3 Evaluation methods

We evaluated the performance of *seq2squiggle* and squigulator through two modes: Read-mode and Genome-mode.

##### Read-mode

Here, signals were simulated based on experimentally basecalled reads (FASTQ/A data). This approach enabled a direct comparison between simulators and the experimental data using the same set of reads, focusing on signal similarity and basecalling metrics. We used dynamic time warping (DTW) to assess signal similarity because it effectively handles variations in signal length and temporal alignment, which are common in nanopore sequencing data. We employed mrmsdtw (Pratzlich *et al*., 2016), a memory-efficient implementation of DTW provided by the linmdtw package (Tralie and Dempsey, 2020). Mrmsdtw offers reduced memory usage compared to the traditional DTW implementation, making it suitable for handling large-scale nanopore signal comparisons. We normalized the mrmsdtw values to account for read length differences and limited our analysis to 2,000 reads to manage computational demands.

##### Genome-mode

In this mode, simulators generated reads from the same input genome (FASTA data). Following this, signals are simulated based on these generated reads. This allowed us to evaluate the simulators’ ability to replicate the quality of real nanopore sequencing reads. Key metrics included successful alignment rate, alignment ratio, match rate, mismatch rate, insertion rate, deletion rate, and the AUC of the average match rate, as described previously (Pagès-Gallego and De Ridder, 2023).

Additionally, we examined the variant detection rate by integrating high-confidence NA12878 variants from Genome in a Bottle (v3.3.2) into chromosome 22 of the human reference genome, similar to previous work by Gamaarachchi and coworkers (Gamaarachchi *et al*., 2023), before undergoing simulation with *seq2squiggle* and squigulator. 25o,000 reads generated by each simulator were basecalled and aligned to the default hg38 reference genome, following the same procedures as previously described. Variant calling was performed using *Clair3* (Zheng *et al*., 2022) and the results were evaluated using *RTGtools* (Cleary *et al*., 2014) against the integrated high-confidence variants. Detailed commands and additional information are provided in the Supplementary Data.

## 3 Results

### 3.1 Signal similarity and basecalling accuracy

To demonstrate the performance of *seq2squiggle*, we conducted experiments on multiple datasets and compared our tool with squigulator, the only available simulator for R10.4.1 chemistry, as well as with experimental data. We selected squigulator because it outperformed DeepSimulator and demonstrated strong performance on R9.4 data (Gamaarachchi *et al*., 2023), making it a suitable baseline for our evaluation. Other simulators, such as DeepSimulator and NanosigSim, were not considered for comparison as they did not support the latest R10.4.1 chemistry.

Initially, we compared both simulators in Read-mode, where they simulated the same set of experimental input reads. For this comparison, we evaluated both tools in their default mode, which added noise to both the event-length and signal-amplitude domains. In Read-mode, we compared the DTW distance of each simulated read to the corresponding real experimental read (Supplementary Figure 1). The results indicated that reads generated by squigulator exhibited a slightly larger DTW deviation (median 9.75) from the real experimental reads compared to those generated by *seq2squiggle* (median 9.58). This represents a small but notable increase in DTW deviation for squigulator. However, while DTW distance provides an initial measure of signal similarity, it does not differentiate between DNA bases and thus does not fully capture the impact on downstream analysis. DTW measures overall signal similarity but cannot account for how deviations in the signal might affect the accuracy of basecalling. For certain DNA bases, these deviations might be tolerable and still result in correct basecalls, whereas for others, they could significantly alter the basecalling outcome. Such nuances are not reflected by DTW alone and can only be understood through downstream analysis.

To evaluate the practical significance of these deviations, we extended our comparison to basecalling accuracy and alignment metrics in both Read-mode and Genome-mode using two publicly available datasets. First, we compared the rate of successfully aligned reads and the alignment ratio (Figure 3A-B). Here, *seq2squiggle’s* reads showed a higher alignment ratio and successful alignments in both Read-mode and Genome-mode. As expected, the alignment ratio in Genome-mode was higher for both simulators since reads were generated directly from the target genome. In Read-mode, however, the signals were generated from basecalled experimental reads, which may contain basecalling errors that impact the alignment accuracy. Our proposed model generated reads with an overall higher median match rate of 91.55 (human Read-mode) to 92.91 (human Genome-mode) across all datasets (Figure 3C). In contrast, squigulator showed a lower match rate of 83.25 to 86.28. Further, *seq2squiggle* showed a lower mismatch rate (Figure 3D), a lower deletion rate, and a lower insertion rate (Supplementary Figure 2) compared to squigulator. Notably, the differences in insertion rates were smaller compared to the differences in mismatch and deletion rates. Reads generated by *seq2squiggle* exhibited an overall higher median PHRED score of 11.42 to 11.82, while squigulator generated reads with a median Q-score of 7.51 to 8.24 (Figure 3E). Finally, we plotted the AUC of the average match rate of the reads sorted by the average PHRED score (Supplementary Figure 3). Again, *seq2squiggle* showed higher AUC values across all datasets compared to squigulator (Figure 3F). We also compared the read length distribution of both tools with the experimental data (Supplementary Figure 2D). Here, both simulators exhibited realistic read length distributions, although variations were observed due to factors such as the species being sequenced, the type of sequencing device used, and other experimental parameters.

**Figure 3.**
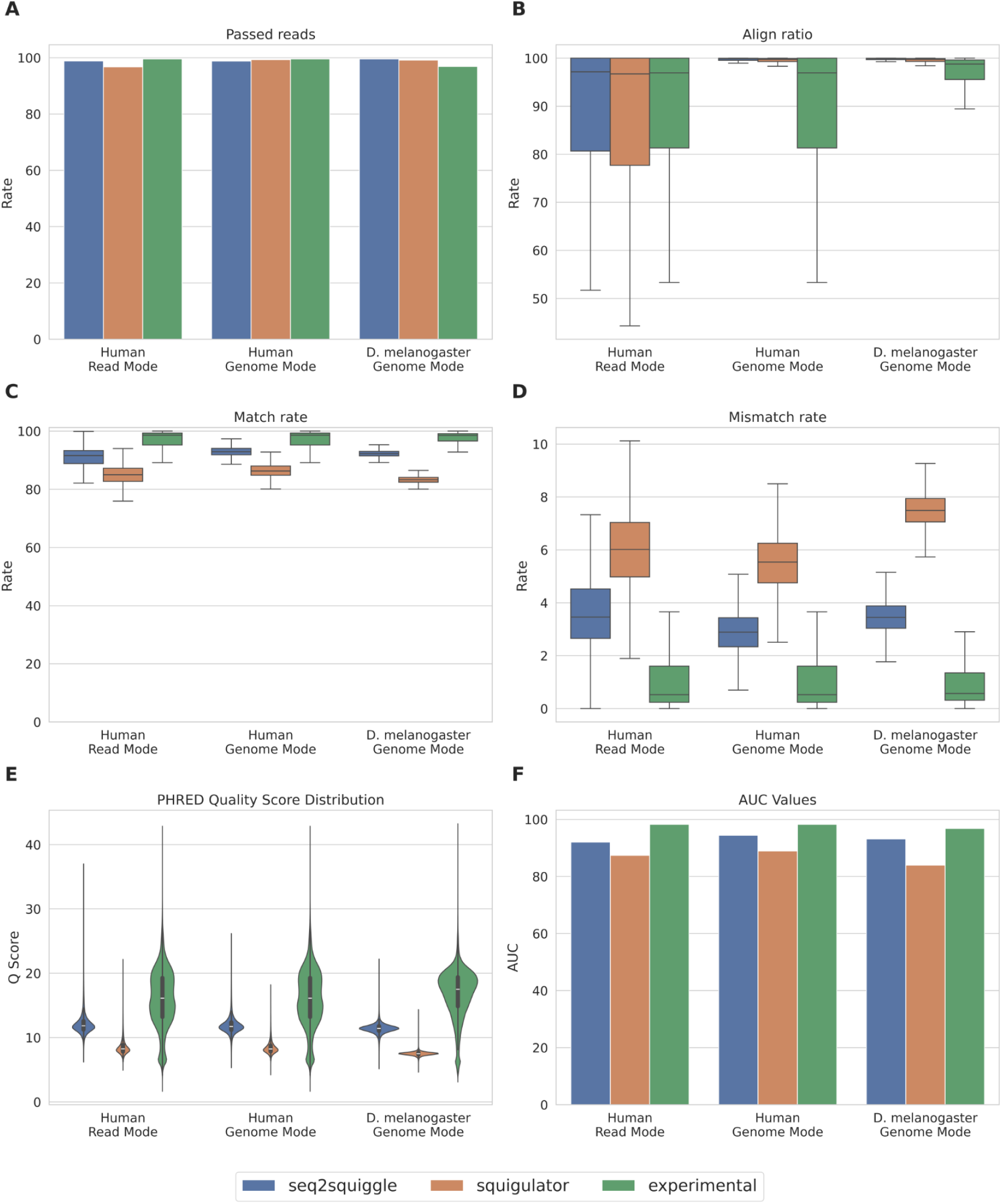
Performance comparison of seq2squiggle (blue), squigulator (orange), and experimental data (green) across multiple datasets and several performance metrics. (A) Proportion of successfully aligned reads to the reference genome. (B) Distribution of aligned bases to the reference genome. (C) Distribution of match rates. (D) Distribution of mismatch rates. (E) Distribution of PHRED quality scores (F) Area under the curve (AUC) values for match rates sorted by average PHRED scores.

### 3.2 Detection of SNVs and Indels

In this section, we present the results of our analysis of single nucleotide variations (SNVs) and insertions/deletions (indels) using simulated data. We simulated 250,000 reads from chromosome 22 of the human reference genome using both squigulator and *seq2squiggle*.

Reads generated with *seq2squiggle* showed a higher number of true-positive SNVs, along with an overall higher recall and precision compared to squigulator (Figure 4). However, upon examining the performance of Indel variants (Supplementary Figure 6-7), both tools displayed relatively high false-positive rates. The challenges in detecting small Indels with nanopore sequencing, attributed to homopolymer-induced errors, were further exacerbated by the insertion and deletion rates observed in both *seq2squiggle* and squigulator (Supplementary Figure 2). Hence, the average match rate in homopolymer regions for *seq2squiggle* and squigulator was generally lower compared to experimental data (Supplementary Figure 8).

**Figure 4.**
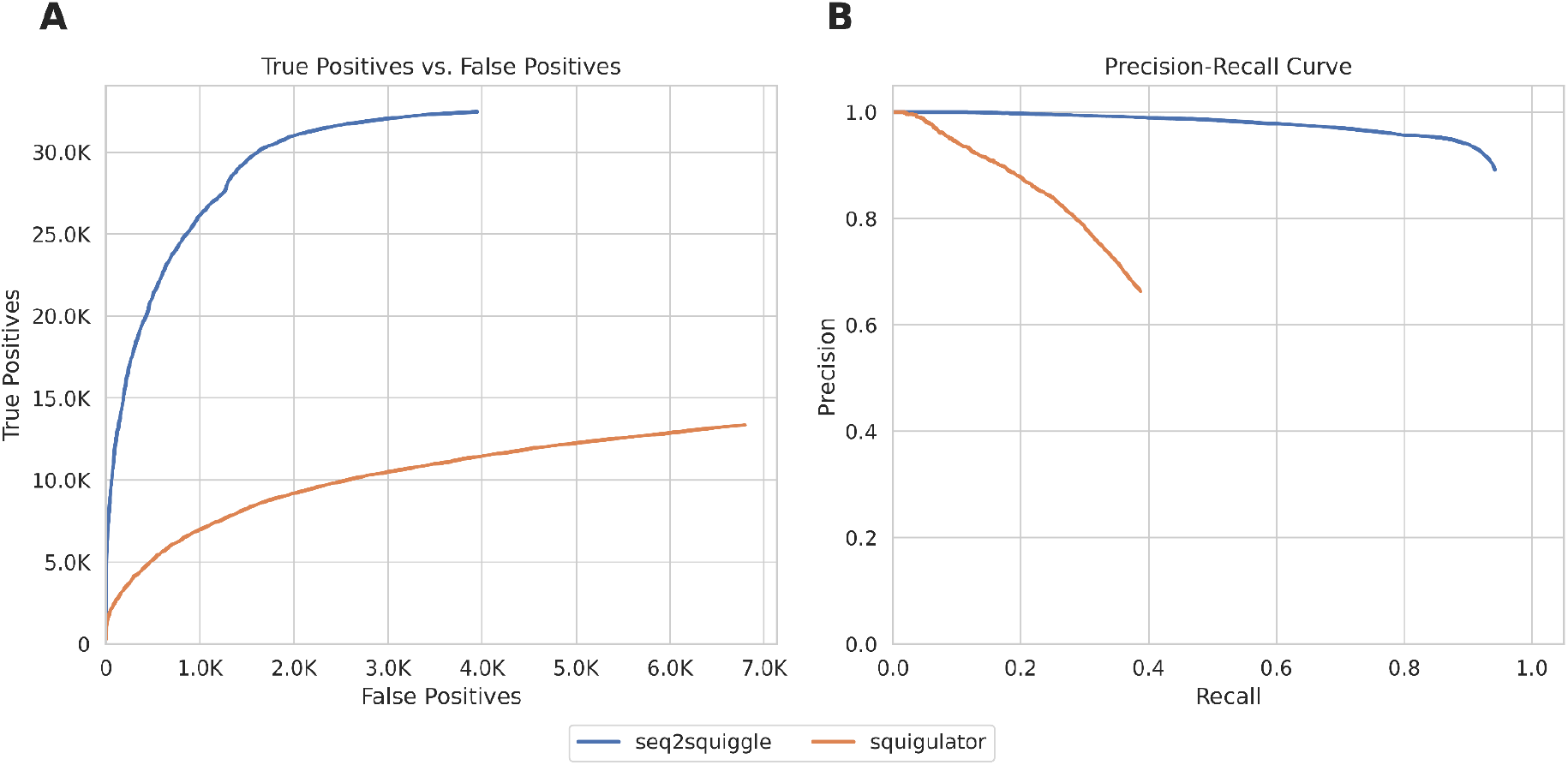
Evaluation of SNP detection accuracy by Clair3 comparing seq2squiggle (blue) and squigulator (orange) using RTGtools’ SNP ROC data. (A) Receiver Operating Characteristic (ROC) curve illustrating the relationship between the total number of false positives and true positives. (B) Precision-Recall curve showing the precision and recall performance.

Furthermore, we inspected the number of false-positive SNV and Indel calls based on genomic regions, specifically homopolymer regions (defined as sequences with at least five consecutive identical bases), tandem repeat regions (defined using Tandem Repeat Finder (Benson, 1999)), and regions without homopolymers or tandem repeat elements (Supplementary Figure 9). Our analysis revealed that homopolymer regions yielded a lower percentage of true positives for both tools. However, *seq2squiggle* demonstrated a higher number of true-positive variant calls across all three regions.

### 3.3 Influence of noise on performance

We examined the impact of noise parameters and modules on the basecalling performance, a topic previously noted by the authors of squigulator who observed differing effects of noise in the time and amplitude domains on basecalling performance. It was found that higher noise levels in event lengths reduced the accuracy of basecalling. However, intriguingly, optimal basecalling performance was not achieved when no additional amplitude noise was introduced. This observation can be attributed to the basecaller’s training on experimental data, which inherently contains noise in the amplitude domain (Gamaarachchi *et al*., 2023). Here, we conducted similar experiments, where we executed *seq2squiggle* and *squigulator* across four noise modes: (i) no noise in the amplitude or event-length domain (ii) noise in the amplitude domain only (iii) noise in the event-length domain only and (iv) noise in both the amplitude and event-length domains. These experiments were performed using a subset of 50,000 reads from the *D. melanogaster* dataset. Consistent with prior research, we observed that the incorporation of noise in the amplitude domain enhanced the average match rate compared to noise-free signal simulation in both *seq2squiggle* and squigulator (Supplementary Figure 4-5). For *seq2squiggle*, the AUC value increased from 81.61 to 93.04, while for squigulator, it increased from 76.39 to 87.44. Introducing additional noise in the event-length domain resulted in a reduced average match rate, indicating that varying event durations did not significantly aid in distinguishing different DNA bases during basecalling. Introducing duration-noise involved balancing the generation of realistic signals with a slight compromise in performance. Comparing the AUC value of *seq2squiggle* and squigulator, we observed that *seq2squiggle* outperformed squigulator across all four noise modes based on the AUC value (Table 2).

**Table 2.** Area under the curve (AUC) values of match rates sorted by PHRED scores for seq2squiggle and squigulator in four different noise modes: (1) no noise in amplitude or event-length domain, (2) Noise in amplitude domain only, (3) Noise in event-length domain only, and (4) Noise in both amplitude and event-length domains.

| Noise in amplitude domain | Noise in event-length domain | AUC values of *seq2squiggle* | AUC values of squigulator |
| --- | --- | --- | --- |
|  |  | 81.61 | 76.39 |
| ✓ |  | 93.04 | 87.41 |
|  | ✓ | 83.48 | 73.09 |
| ✓ | ✓ | 92.94 | 83.83 |

Furthermore, we modified *seq2squiggle* by replacing both the duration sampler and the noise sampler with static normal distributions (mean=9.0, std=4.0 for length, and mean=0.0, std=1.0 for noise), akin to the noise sampling method used in squigulator. This adjustment allowed us to compare the benefits of using duration and noise modules that learned distributions based on DNA sequences, rather than relying on predefined statistical models. Our findings indicated that the noise module in *seq2squiggle* generated signals with a slightly higher match rate (Supplementary Table 3), suggesting that our approach better captured noise patterns that resembled real experimental data. However, the learned duration module resulted in a slightly lower match rate, highlighting that learned variations in event lengths might not enhance performance as effectively as amplitude noise.

## 4 Discussion

In this study, we introduced *seq2squiggle*, an innovative simulator designed for nanopore sequencing data. Unlike existing simulators, our model leverages a feed-forward transformer architecture to learn the intricate relationship between the input DNA sequence and the corresponding nanopore signal sequence directly from the training data. This approach avoids the direct need for pre-defined statistical models and pore models, which are commonly used in other simulators. By avoiding these models, *seq2squiggle* mitigates potential biases and inaccuracies that can arise from relying on static statistical assumptions.

Our experiments have demonstrated that *seq2squiggle* surpassed existing simulators in generating signals that closely resemble real nanopore sequencing data, achieving higher match rates, PHRED scores, SNP detection rates, and other critical performance metrics. Despite these advancements, discrepancies between simulated and experimental data still persist, possibly due to sub-optimal segmentation and the higher complexity of R10.4.1 data. In future work, we aim to improve the architecture of *seq2squiggle* to further increase the basecalling accuracy of simulated reads, aiming for closer alignment with real-world data. Although we currently employ modules in the amplitude and event-length domains to sample noise, we make certain assumptions about the noise distributions, such as normal and gamma distributions. Recent innovations in speech synthesis (Huang *et al*., 2022; Zhang *et al*., 2023), particularly the integration of diffusion modules, show promise for advancing our model. By incorporating more sophisticated noise modeling techniques and leveraging diffusion-based approaches, we anticipate further improvements in the quality of our simulated nanopore sequencing signals. Additionally, we intend to explore the use of *seq2squiggle* for detecting DNA methylations and simulating RNA data, thereby expanding its applicability and utility. Further, the model can be simply retrained to adapt to forthcoming updates to the flow cell chemistry.

It is important to note that *seq2squiggle* has higher runtime and memory consumption compared to squigulator. This is due to its deep learning-based approach, which, while resource-intensive, results in significantly higher quality of the generated signals. The trade-off between computational resources and accuracy is a common challenge in deep learning applications, but the improved performance of *seq2squiggle* justifies its use for applications requiring realistic nanopore signals. Further, its non-autoregressive architecture keeps the runtime in a manageable timeframe to conduct large-scale simulations.

In summary, *seq2squiggle* represents a significant advancement in the simulation of nanopore sequencing data and provides a powerful tool for the nanopore community. Its ability to generate realistic nanopore signals for the latest flow cell generation has great potential for a variety of applications, including method development, benchmarking, and comprehensive genomic analysis.

## Data availability

The benchmarking code and results can be accessed via Figshare (doi.org/10.6084/m9.figshare.26539696). The benchmarking workflow is available at GitHub (https://github.com/ZKI-PH-ImageAnalysis/seq2squiggle-benchmark).

## References

1 Ahsan, M.U. et al. (2024) A signal processing and deep learning framework for methylation detection using Oxford Nanopore sequencing. Nat. Commun., 15, 1448.

2 Amarasinghe, S.L. et al. (2020) Opportunities and challenges in long-read sequencing data analysis. Genome Biol., 21, 30.

3 Baker, E.A.G. et al. (2016) SiLiCO: A Simulator of Long Read Sequencing in PacBio and Oxford Nanopore Genomics. biorxiv. https://www.biorxiv.org/content/10.1101/076901v1 12 August 2024, date last accessed.

4 Benson, G. (1999) Tandem repeats finder: a program to analyze DNA sequences. Nucleic Acids Res., 27, 573–580.

5 Chen, W. et al. (2020) Simulation of Nanopore Sequencing Signals Based on BiGRU. Sensors, 20, 7244.

6 Cleary, J.G. et al. (2014) Joint Variant and De Novo Mutation Identification on Pedigrees from High-Throughput Sequencing Data. J. Comput. Biol., 21, 405–419.

7 Delahaye, C. and Nicolas, J. (2021) Sequencing DNA with nanopores: Troubles and biases. PLOS ONE, 16, e0257521.

8 Gamaarachchi, H. et al. (2022) Fast nanopore sequencing data analysis with SLOW5. Nat. Biotechnol., 40, 1026–1029.

9 Gamaarachchi, H. et al. (2023) Squigulator: simulation of nanopore sequencing signal data with tunable noise parameters. biorxiv. https://www.biorxiv.org/content/10.1101/2023.05.09.539953v1 12 August 2024, date last accessed.

10 Giesselmann, P. (2021) Genome Analysis Methods using Long Read Nanopore Sequencing. 10.17169/refubium-32662 12 August 2024, date last accessed.

11 Huang, R. et al. (2022) ProDiff: Progressive Fast Diffusion Model For High-Quality Text-to-Speech. arxiv. https://arxiv.org/abs/2207.06389 12 August 2024, date last accessed.

12 Jenkins, V.K. et al. (2022) Using FlyBase: A Database of Drosophila Genes and Genetics. In, Dahmann, C. (ed), Drosophila, Methods in Molecular Biology. Springer US, New York, NY, pp. 1–34.

13 Kovaka, S. et al. (2024) Uncalled4 improves nanopore DNA and RNA modification detection via fast and accurate signal alignment. biorxiv. https://www.biorxiv.org/content/10.1101/2024.03.05.583511v1 12 August 2024, date last accessed.

14 Li, Y. et al. (2018) DeepSimulator: a deep simulator for Nanopore sequencing. Bioinformatics, 34, 2899–2908.

15 Liang, S. and Zhang, Y. (2020) A Simple General Approach to Balance Task Difficulty in Multi-Task Learning. arxiv. https://arxiv.org/abs/2002.04792 12 August 2024, date last accessed.

16 Pagès-Gallego, M. and De Ridder, J. (2023) Comprehensive benchmark and architectural analysis of deep learning models for nanopore sequencing basecalling. Genome Biol., 24, 71.

17 Pratzlich, T. et al. (2016) Memory-restricted multiscale dynamic time warping. In, 2016 IEEE International Conference on Acoustics, Speech and Signal Processing (ICASSP). IEEE, Shanghai, pp. 569–573.

18 Ren, Y. et al. (2022) FastSpeech 2: Fast and High-Quality End-to-End Text to Speech. arxiv. https://arxiv.org/abs/2006.04558 12 August 2024, date last accessed.

19 Ren, Y. et al. (2019) FastSpeech: Fast, Robust and Controllable Text to Speech. arxiv. https://arxiv.org/abs/1905.09263 12 August 2024, date last accessed.

20 Samarakoon, H. et al. (2023) Accelerated nanopore basecalling with SLOW5 data format. Bioinformatics, 39, btad352.

21 Teng, H. et al. (2018) Chiron: translating nanopore raw signal directly into nucleotide sequence using deep learning. GigaScience, 7, giy037.

22 Tralie, C. and Dempsey, E. (2020) Exact, Parallelizable Dynamic Time Warping Alignment with Linear Memory. arxiv. https://arxiv.org/abs/2008.02734 12 August 2024, date last accessed.

23 Wang, Yunhao et al. (2021) Nanopore sequencing technology, bioinformatics and applications. Nat. Biotechnol., 39, 1348–1365.

24 Zhang, Chenshuang et al. (2023) A Survey on Audio Diffusion Models: Text To Speech Synthesis and Enhancement in Generative AI. arxiv. https://arxiv.org/abs/2303.13336 12 August 2024, date last accessed

25 Zheng, Z. et al. (2022) Symphonizing pileup and full-alignment for deep learning-based long-read variant calling. Nat. Comput. Sci., 2, 797–803.

