## Supplementary Data for "End-to-end simulation of nanopore sequencing signals with feed-forward transformers"

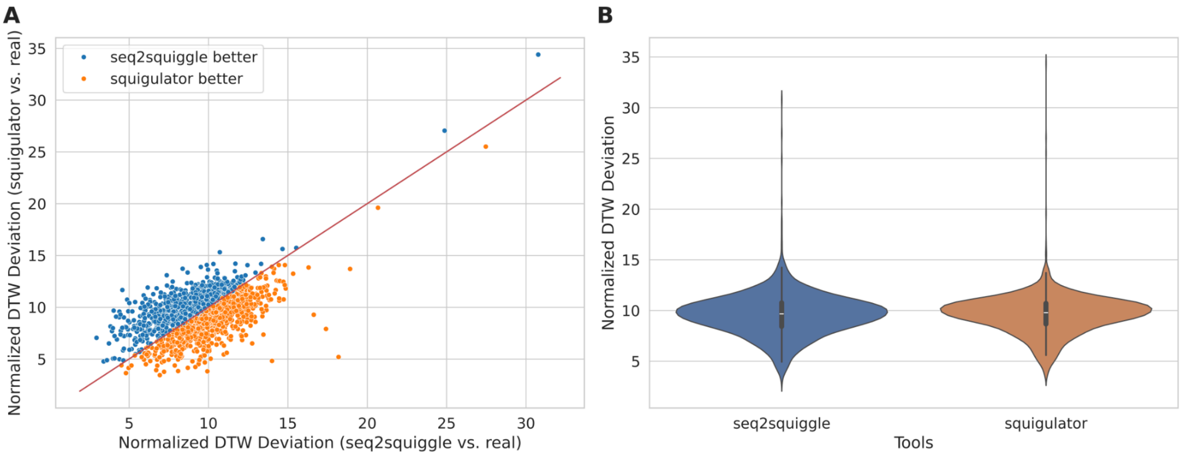


Supplementary Figure 1. Comparison of DTW deviations between the simulators and the experimental data in human read mode. (A) Scatterplot illustrating the DTW deviation between simulated and real signals. Each point represents a read, displaying the DTW deviation of seq2squiggle (X-axis) and squigulator (Y-axis) relative to the real data. Points above the red diagonal line indicate *seq2squiggle* exhibits a smaller DTW deviation compared to squigulator, and vice versa. (B) Violin plot depicting the distribution of DTW deviations between both simulators and real data.

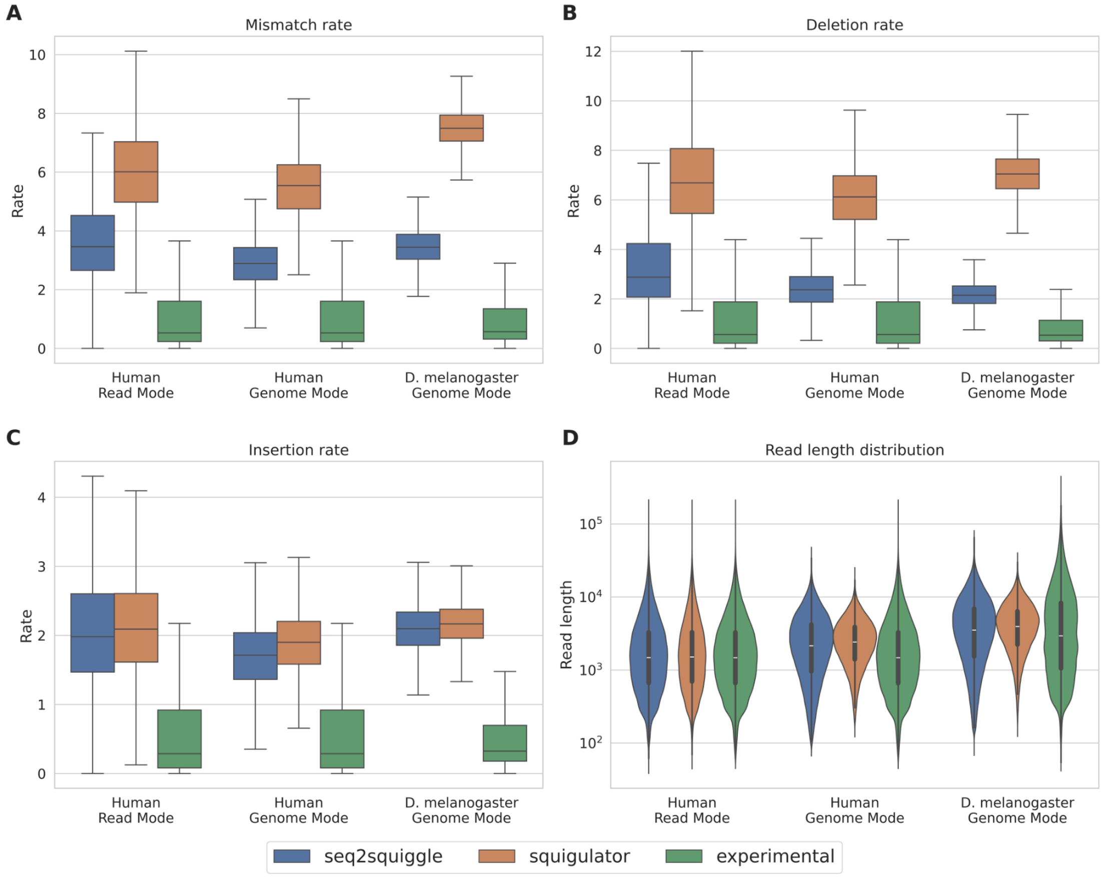


Supplementary Figure 2. Performance comparison of seq2squiggle (blue), squigulator (orange), and experimental data (green) across multiple datasets and several performance metrics. (A) Distribution of mismatch rates. (B) Distribution of deletion rates. (C) Distribution of insertion rates. (D) Distribution of read lengths.

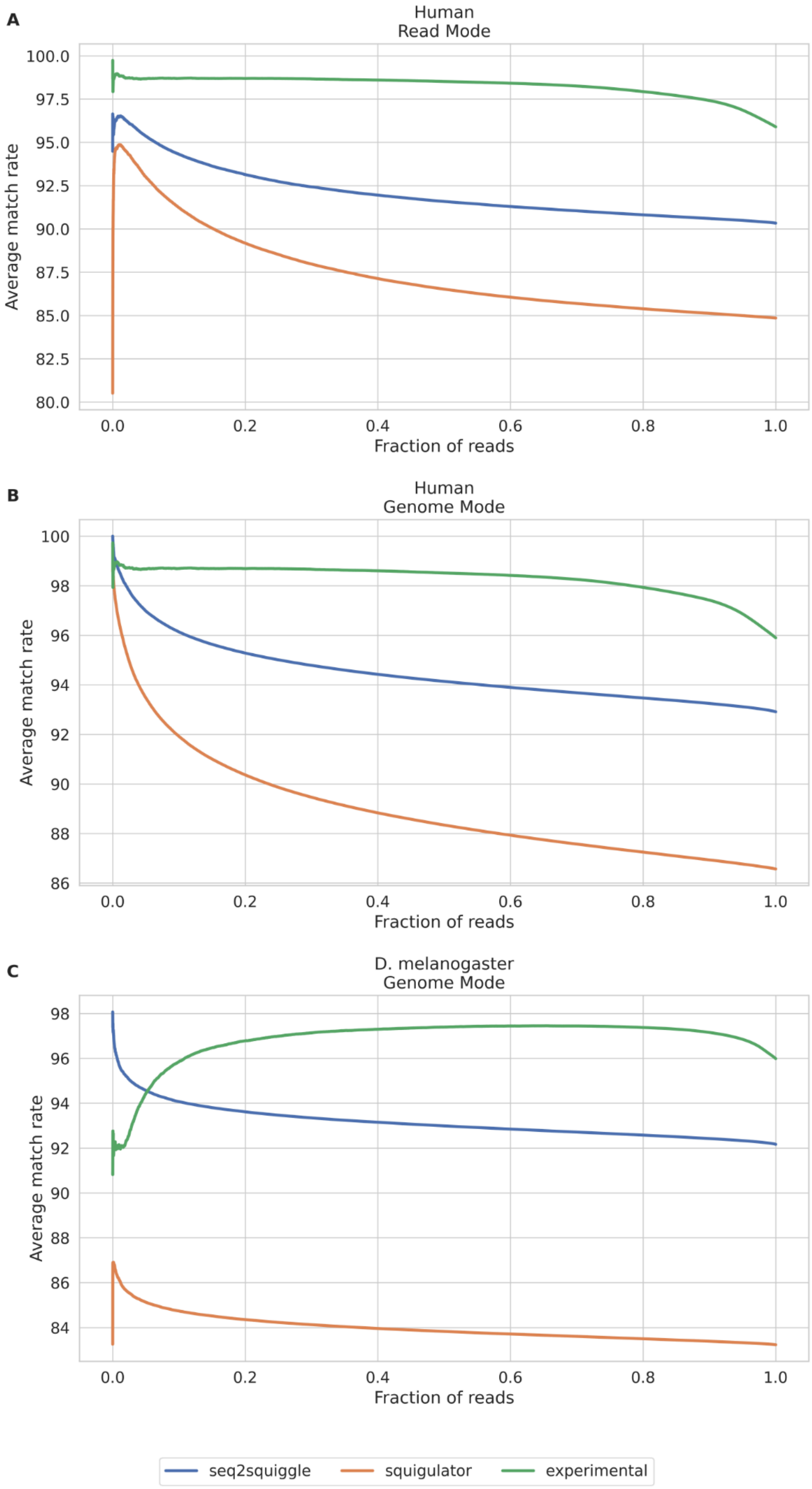


Supplementary Figure 3. AUC of match rate sorted by average PHRED score for seq2squiggle (blue), squigulator (orange), and experimental data (green) across multiple datasets.

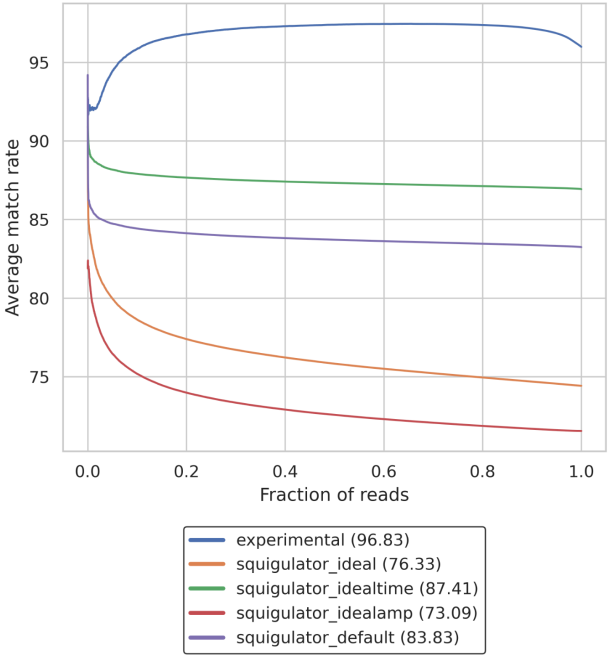


Supplementary Figure 4. AUC of match rate sorted by average PHRED score using different noise modes for squigulator.

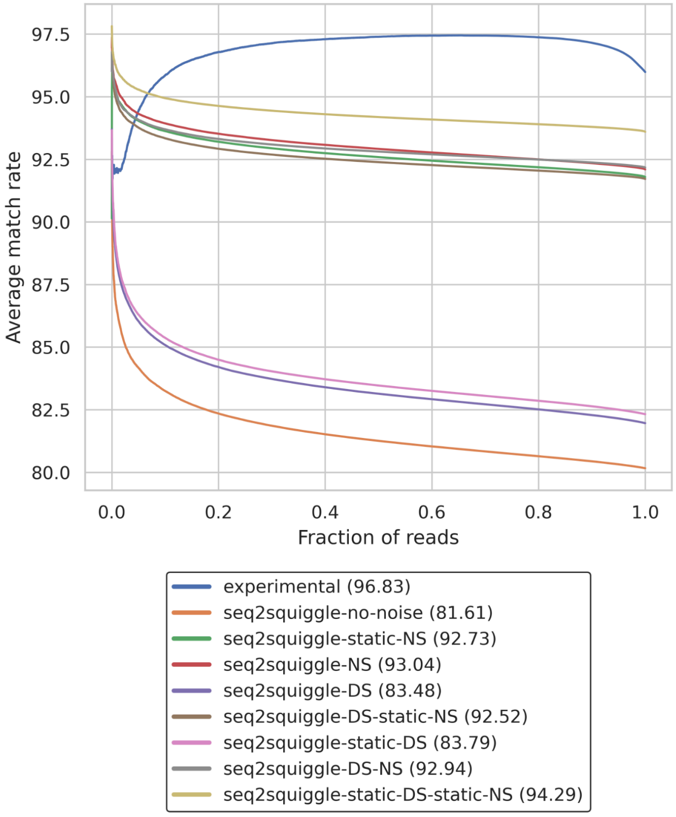


Supplementary Figure 5. AUC of match rate sorted by average PHRED score using various noise modes for *seq2squiggle*. The noise modes include manual Noise Sampling with static distribution (static NS), Noise Sampling with Noise Sampler (NS), manual duration sampling with static distribution (static DS) and duration sampling with Duration Sampler (DS).

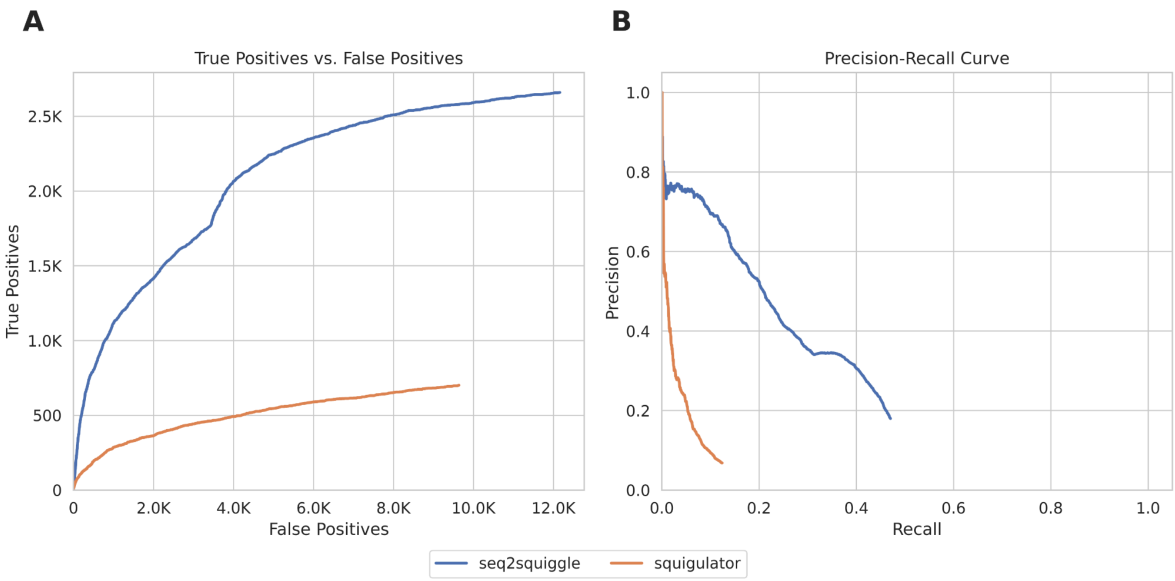


Supplementary Figure 6. Accuracy evaluation of SNP and Indel detection by Clair3 comparing squigulator and seq2squiggle using RTGtools' weighted ROC data. (A) Receiver Operating Characteristic (ROC) curve illustrating the relationship between the total number of false positives and true positives. (B) Precision-Recall curve showing the precision and recall performance.

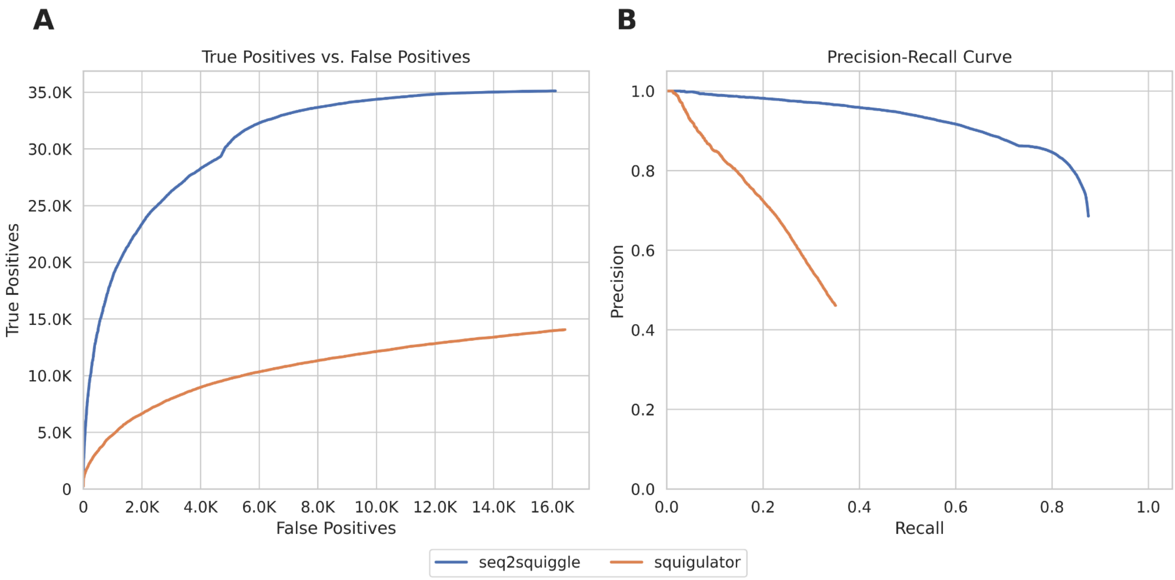


Supplementary Figure 7. Accuracy evaluation of Indel detection by Clair3 comparing squigulator and seq2squiggle using RTGtools' Indel ROC data. (A) Receiver Operating Characteristic (ROC) curve illustrating the relationship between the total number of false positives and true positives. (B) Precision-Recall curve showing the precision and recall performance.

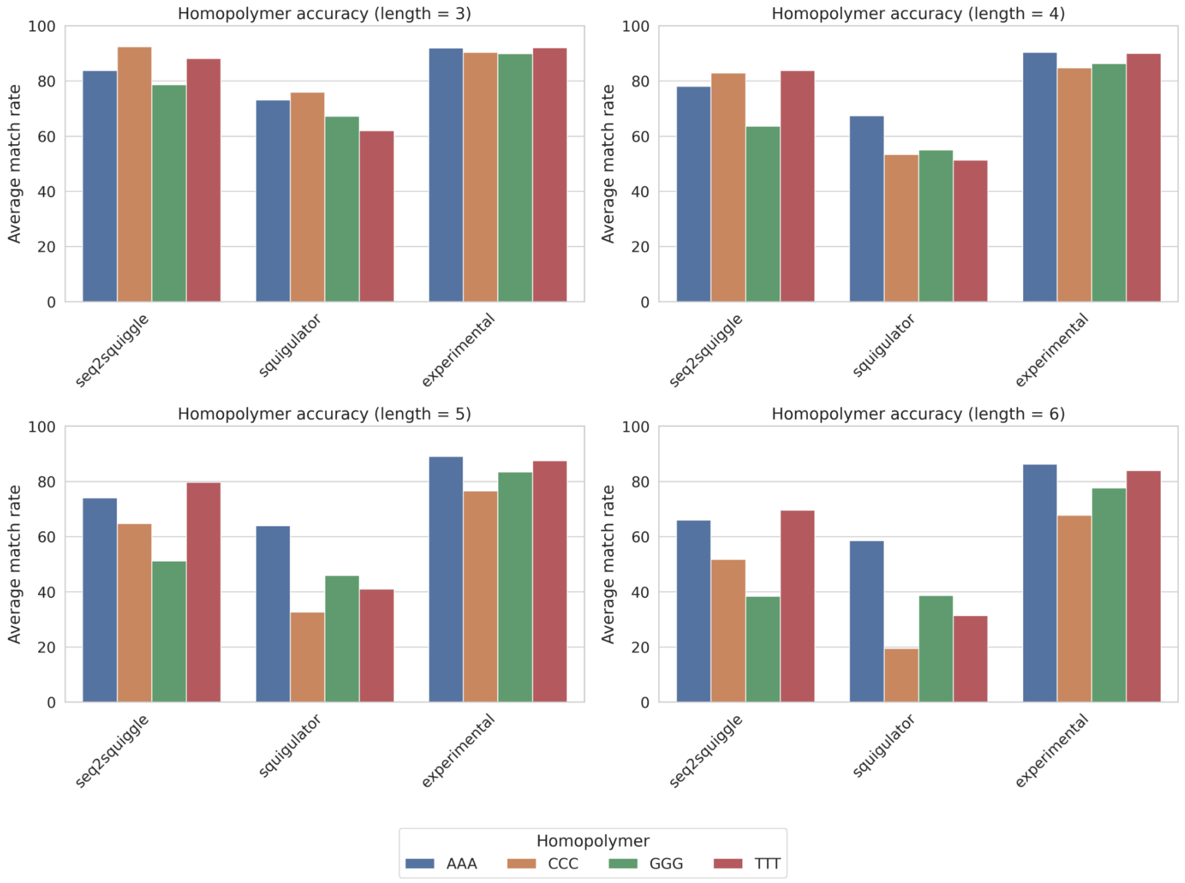


Supplementary Figure 8. Average match rate for different homopolymer lengths on the human dataset. Squigulator and seq2squiggle were executed in genome mode.

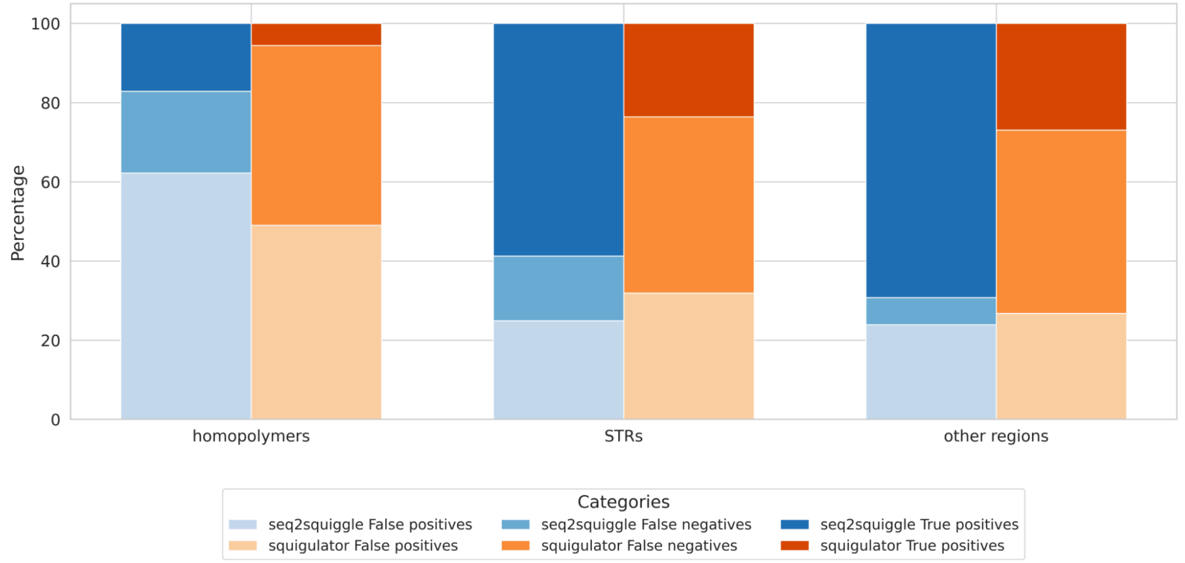


Supplementary Figure 9. Distribution of false positives, false negatives, and true positives across various genomic regions. The relative proportions of these metrics are presented for squigulator (orange) and seq2squiggle (blue) in homopolymer regions, short tandem repeat regions, and other genomic areas. Homopolymer regions are defined as sequences with at least five consecutive identical bases, while short tandem repeat regions are identified using Tandem Repeats Finder.

Supplementary Table 1 - Model hyperparameter for seq2squiggle

| Parameter | Value |
|---|---|
| DNA Embedding dimension | 64 |
| Pre-Net Layers | 1 |
| Pre-Net Hidden | 64 |
| Encoder FFT Layers | 2 |
| Encoder FFT Hidden | 64 |
| Encoder Attention Heads | 8 |
| Encoder feed forward upwards projection size | 256 |
| Decoder FFT Layer | 2 |
| Decoder FFT Hidden | 64 |
| Decoder Attention Heads | 8 |
| Decoder feed forward upwards projection size | 256 |
| Duration Sampler Hidden | 64 |
| Noise Sampler Hidden | 64 |
| Dropout | 0.1 |
| Batchsize | 512 |
| Learning Rate | 0.00025 |
| Warmup Ratio | 0.01 |
| Learning Rate schedule | Linear warmup followed by cosine decay |
| Gradient clipping value | 1.0 |
| Optimizer | Adam |
| Total number of parameters | 219,780 |

Supplementary Table 2 - Average runtime and memory usage of *seq2squiggle* and squigulator for generating 100,000 human reads in Genome-mode. Both tools were run with 64 threads, with *seq2squiggle* using a single A100 GPU.

| Tool | seq2squiggle | squigulator |
|---|---|---|
| Runtime (in hour, minutes, seconds format) | 0:21:15 | 0:01:15 |
| max_rss (in MegaByte) | 121947.91 | 3175.67 |
| max_vms (in MegaByte) | 3305409.93 | 3660.41 |
| max_uss (in MegaByte) | 6839.41 | 3174.10 |
| max_pss (in MegaByte) | 7340.03 | 3174.20 |
| io_in (in MegaByte) | 257.61 | 3000.99 |
| in_out (in MegaByte) | 9827.29 | 6994.70 |
| mean_load | 200.77 | 353.72 |
| cpu_time | 2580.10 | 268.57 |

Supplementary Table 3 - Area Under the Curve (AUC) of match rate sorted by PHRED score for the default implementation *seq2squiggle* using learned noise modules and *seq2squiggle* using static normal distribution for noise (mean=9.0, std=4.0) and event-length (mean=0.0, std=1.0) sampling. The comparison is made across four different noise modes.

| Noise in amplitude domain | Noise in event-length domain | learned noise modules | static noise modules |
|---|---|---|---|
| ✓ | | 93.04 | 92.73 |
| | ✓ | 83.48 | 83.79 |
| ✓ | ✓ | 92.94 | 94.29 |
